# Metabolite profile and mitochondrial energetics characterize poor early recovery of muscle mass following hind limb unloading in old mice

**DOI:** 10.1101/183244

**Authors:** Xiaolei Zhang, Michelle B. Trevino, Miao Wang, Stephen J. Gardell, Julio E. Ayala, Xianlin Han, Daniel P. Kelly, Bret H. Goodpaster, Rick B. Vega, Paul M. Coen

## Abstract

The progression of age-related sarcopenia can be accelerated by impaired recovery of muscle mass following periods of disuse due to illness or immobilization. The molecular underpinnings of poor recovery of aging muscle following disuse remain largely unknown. However, recent evidence suggests that mitochondrial energetics may play an important role. Here, we report that 22-24 month old mice with low muscle mass and insulin resistance display poor early recovery of muscle mass following 10 days of hind limb unloading. We took an unbiased approach to identify changes in energy metabolism gene expression and metabolite pools and show for the first time that persistent mitochondrial dysfunction, dysregulated fatty acid β-oxidation and elevated H_2_O_2_ emission underlie poor early recovery of muscle mass following a period of disuse in old mice. Importantly, this is linked to more severe whole-body insulin resistance. The findings suggest that muscle fuel metabolism and mitochondrial energetics should be a focus for mining therapeutic targets to improve recovery of muscle mass following periods of disuse in older animals.

## INTRODUCTION

Skeletal muscle is a highly plastic tissue that can adapt its mass, structure and metabolic capacity in response to changes in mechanical load. During aging, skeletal muscle exhibits a progressive loss of mass and strength, also called sarcopenia, and an attenuated adaptive recovery following atrophy induced by: disuse, immobilization, starvation and glucocorticoid treatment (Dardevet *et al*. 1995; Zarzhevsky *et al*. 2001; Degens & Alway 2003; Hao *et al*. 2011; Magne *et al*. 2011). The hypertrophic response to functional overload is also blunted with aging (Degens & Alway 2003). Poor recovery of muscle mass and strength, even with exercise training, a potent anabolic stimuli, has also been reported in older humans following unilateral leg immobilization (Suetta *et al*. 2013). Indeed, the lack of recovery from disuse has been suggested to accelerate sarcopenia in older adults, a condition that can increase risk of frailty, morbidity and loss of independence (English & Paddon-Jones 2010). Currently, there are no effective therapies to counteract disuse-induced muscle atrophy and poor recovery of muscle mass, in part because a detailed understanding of the mechanisms underlying loss of muscle plasticity with aging are unknown. In addition, the older population demographic is growing and tends to experience more frequent and prolonged periods of inactivity, increasing the need for the development of safe and effective therapeutics.

Maintenance of skeletal muscle mass reflects a finely controlled balance between protein synthesis and degradation. Hypertrophy results from a shift to protein synthesis and the primary anabolic pathway involves the mammalian target of rapamycin complex 1 (mTORC1). mTORC1 is principally regulated by nutritional cues, insulin/growth factor signaling cascades and mechanotransduction (Hoppeler 2016). Contractile inactivity induced muscle atrophy is due to reduced basal and postprandial protein synthesis, also known as anabolic resistance, and elevated protein degradation by ubiquitin/proteasome and autophagy/lysosome pathways (Bonaldo & Sandri 2013). Protein breakdown is initiated by inflammatory, glucocorticoid, and myostatin/TGFβ pathways (Egerman & Glass 2014). While the protein degradation/synthesis pathways are relatively well characterized, upstream regulation, particularly in the context of muscle plasticity in aging, is less well understood.

Separate lines of investigation have shown that mitochondrial dysfunction occurs in aging muscle, as evidenced by decreased content, mitochondrial protein synthesis, elevated reactive oxygen species (ROS) emission and apoptosis, and reduced ATP production, calcium handling (Joseph *et al*. 2015). Recent evidence also points to mitochondria as an important nexus of control for acute muscle atrophy during disuse (Powers *et al*. 2012). Rodent studies using mitochondrial-targeted antioxidants support a crucial role for mitochondrial ROS in mediating muscle atrophy, through upregulation of ubiquitin-protease enzymes (Min *et al*. 2011; Talbert *et al*. 2013). In addition, a model of increased ROS production, the superoxide dismutase (Sod1) knockout, shows exacerbated muscle atrophy during aging (Jang *et al*. 2010). Elevated mitochondrial oxidative stress is purported to stimulate muscle protein breakdown by activating lysosome-autophagy and ubiquitin-proteasome systems (Grune *et al*. 2003; Li *et al*. 2003; Aucello *et al*. 2009; Smuder *et al*. 2010) and energetic stress (reduced ATP production) may also activate the AMP kinase (AMPK)-FoxO3 pathways leading to increased protein degradation (Greer *et al*. 2007; Romanello *et al*. 2010). Moreover, over-expression of PGC-1α, a key driver of mitochondrial biogenesis, can protect muscle mass from acute atrophy due to immobilization or disuse (Sandri *et al*. 2006; Brault *et al*. 2010; Cannavino *et al*. 2014). Key regulators of autophagy and mitochondrial fission also regulate muscle mass (Masiero & Sandri 2010; Romanello *et al*. 2010).

Taken together, this body of evidence implicates a central role for mitochondrial energetics in regulating muscle mass and likely plays a vital role in recovery of muscle mass following a period of disuse. The goal of this study was to determine whether age-associated alterations in muscle mass (sarcopenia), mitochondrial function and gene/metabolic profile are linked with responses to hind limb unloading induced atrophy and recovery during reloading in adult (6 mo) and old mice (22-24 mo). Our data suggest that dysregulated fatty acid β-oxidation (FAO) and elevated H_2_O_2_ emission from mitochondria likely contribute to compromised recovery of soleus muscle mass following unloading in old sarcopenic mice.

## RESULTS

### Old mice are resistant to recovery of muscle mass following unloading induced muscle Atrophy

The overarching goal of the following experiments was to determine whether alterations in fuel metabolism and mitochondrial energetics are linked to poor recovery of muscle mass in old mice (22-24 mo), compared to adult mice (6 mo). To assess differences in the atrophy recovery response of old mice, we used a modified version of the tail suspension hind limb unloading model to induce hind limb muscle atrophy over a 10-day time course (Morey-Holton & Globus 2002) as shown in Figure 1A, followed by 3 days of reloading and voluntary cage ambulation. Body weight was greater in the old mice and decreased slightly during unloading and reloading in adult and old mice (Figure 1B). There was no significant change in daily food intake during unloading and reloading in adult and old mice (Figure 1C). As shown in Fig. 1D, E & F, hind limb muscle groups in the old mice weighted less compared to young mice consistent with a sarcopenic phenotype. While it is clear from previous reports that a full recovery of muscle mass and strength may take up to 40 days after unloading (Magne *et al*. 2011), we examined the response following a reloading period of 3 days, a time that others have shown to be important for induction of anabolic signal transduction and gene transcription (Akt, p70S6K, 4E-BP1, STAT3) involved in hypertrophy and recovery of muscle mass (Childs *et al*. 2003; Baehr *et al*. 2016). Our goal was to understand the differences in response between adult and old mice at this key early period during reloading. Unloading induced significant atrophy of the soleus, gastrocnemius and quadriceps muscle groups in both adult and old animals. The soleus is a postural muscle consisting predominantly of type IIA and I oxidative myofibers making it particularly susceptible to unloading induced atrophy. For this reason, we focused our analysis predominantly on the soleus. The magnitude of soleus muscle loss was similar between adult and old mice. Following 3 days of reloading, significant recovery of soleus mass occurred in the adult, but not the old mice (Figure 1D). Similar responses were observed for the gastrocnemius and quadriceps (Figure 1E and 1F).

**Figure 1.**
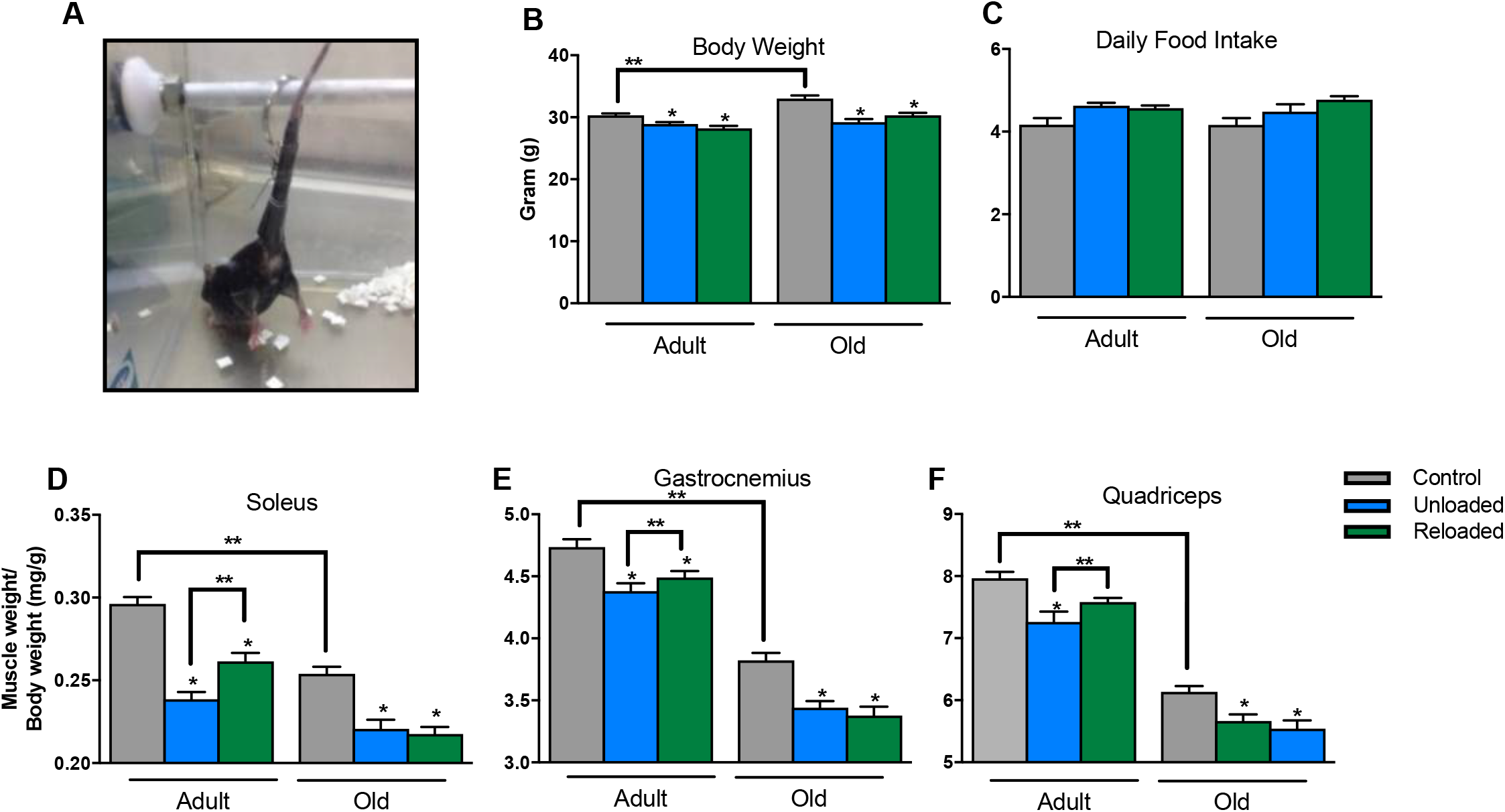
Old mice do not recover muscle mass following 10 days of hind limb unloading and 3 days of recovery, compared to adult mice. <u>Panel A</u>; image of the hind limb unloading approach. <u>Panel B</u>; body weight was greater for the old mice and both adult and old mice lost weight during the unloading and reloading periods. <u>Panel C</u>, average daily food intake was similar for both old and adult mice and did not change over the unloading and reloading periods. <u>Panels D, E and F</u>; Soleus, Gastrocnemius and Quadriceps atrophy during unloading in both adult and old mice. However, after 3 days of reloading, muscle mass did not recover in the old mice. n = 11 per group. Data are presented as mean ± SEM. ** P <0.05 ANOVA/Bonferoni Correction. * P <0.05 v Control ANOVA/Bonferoni Correction.

### Whole body insulin action remains suppressed in old mice following unloading and Recovery

Periods of disuse can induce profound whole body and skeletal muscle insulin resistance. We next examined whether differences in metabolic phenotype may link with the poor recovery of muscle observed in old mice (Figure 2). For this experiment, we compared control animals to reloaded animals. As expected, whole body insulin action (slope of blood glucose response to ITT) was lower in the old compared to adult mice (ANOVA main effect for age, Figure 2A & B). This was likely due to soleus and gastrocnemius glucose uptake being lower in the old compared to adult mice (ANOVA main effect for age, Figure 2C & D). Significantly, we found that whole-body insulin action following reloading was lower in old mice, when compared to control. This was not the case for adult mice (Figure 2A & B). Insulin stimulated muscle glucose clearance was similar between control and reloading in both adult and old mice. However, following reloading, gastrocnemius glucose uptake was significantly greater in adult compared to old mice which likely contributed to lower whole-body insulin action. Body composition as assessed by NMR revealed that old mice had lower lean mass and greater fat mass compared to adult mice (Figure 2E & F, main effect for age), consistent with a sarcopenic phenotype. Following unloading/reloading, old mice had greater fat mass and lower lean mass, changes that again likely contributed to blunted whole body insulin action, and are consistent with the lack of recovery of individual hind limb muscle mass.

**Figure 2.**
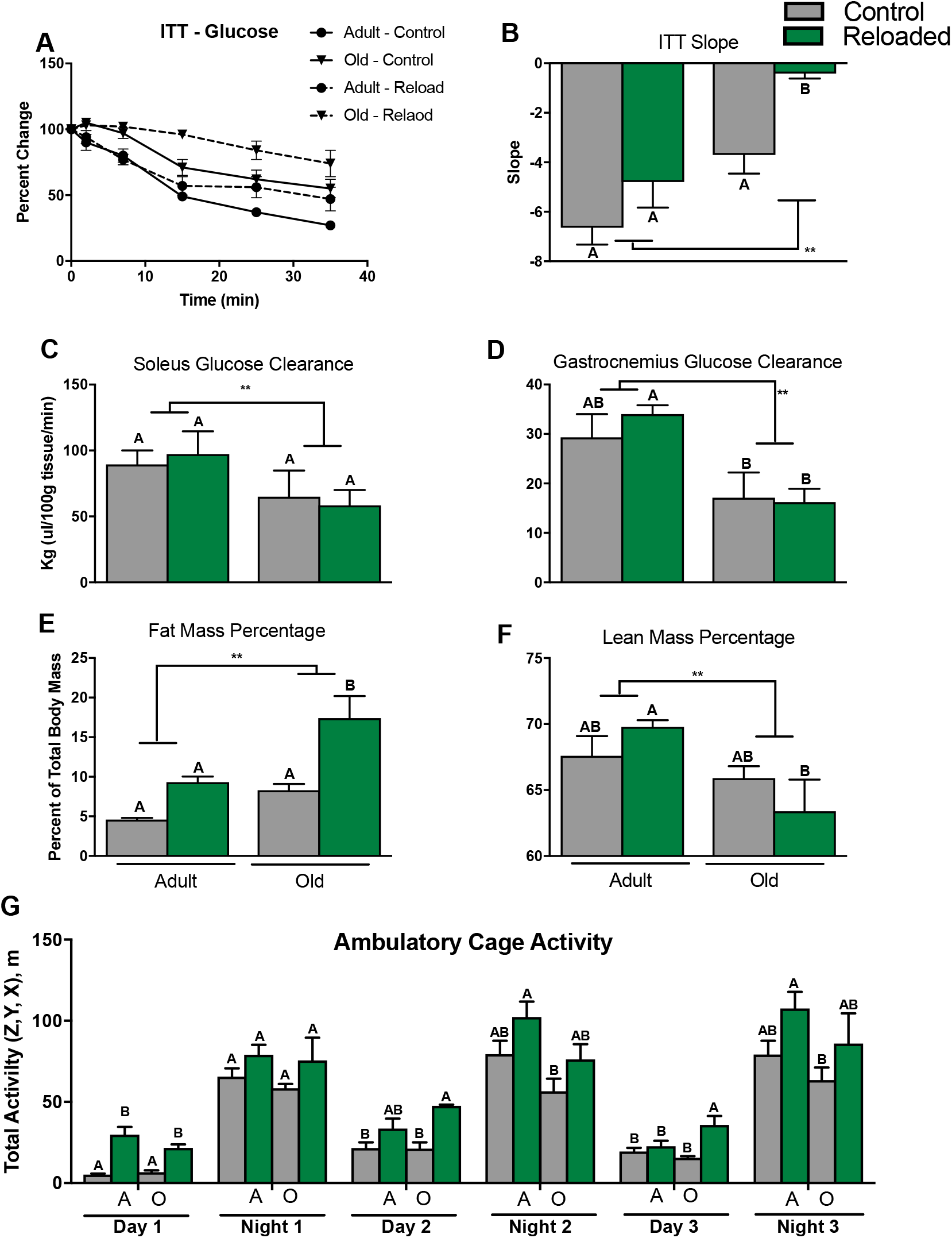
Old mice are insulin resistant and have lower muscle glucose clearance, a phenotype that is exacerbated by unloading and reloading. Grey bars represent control conditions. Green bars represent animals that had 10 days of unloading and 5 days of recovery. <u>Panel A</u>, blood glucose reductions (percent change from baseline) following intraperitoneal insulin injection. <u>Panel B</u>, The slope of the fall in glucose levels from t = 0 to 15 min was used as an index of whole-body insulin action. <u>Panel C</u>, Soleus glucose clearance. <u>Panel D</u>, Gastrocnemius glucose clearance. <u>Panel E</u>, proportion of body fat mass determined by NMR. <u>Panel F</u>, proportion of body lean mass determined by NMR. <u>Panel G</u>, day and night cage ambulatory activity (x,y,z) using the Promethion Mouse Multiplexed Metabolic System. n = 5 per group. Data are presented as mean ± SEM. The letters A and B denote significant differences between group/time points (P < 0.05, ANOVA/Tukey Correction).

Muscle protein synthesis and insulin sensitivity are both very sensitive to muscle contractile activity. To determine whether differences in ambulatory cage activity explained differences in muscle recovery and insulin sensitivity between adult and old mice, we examined cage activity over 3-days of the recovery period (Figure 2G). Generally, both the adult and old mice similarly recovered night and day total ambulatory cage activity, albeit in some cases (Day 1, for example) the reloaded animals were more active compared to controls.

### Inhibition of the AKT/mTOR/FOXO3a pathway in hind-limb unloaded old mice was not rescued by reloading

Maintenance of skeletal muscle mass is a finely controlled balance between protein synthesis and degradation. We next examined the basal phosphorylation status of key mediators of protein synthesis and degradation in the soleus. The AKT/mTOR pathway was inhibited following unloading in both adult and old mice as evidenced by reduced serine phosphorylation of AKT (Ser473), mTOR (Ser2448) and S6 (Ser240/244). Strikingly, inhibition of the AKT/mTOR pathway was rescued by reloading in the adult mice, but not in the old mice (Figure 3A). Forkhead box O3 (FOXO3a) is a transcription factor that regulates the muscle atrophy program. Under catabolic conditions, inhibition of AKT results in dephosphorylation and nuclear translocation of FOXO3a, which activates the expression of Atrogin-1. Consistent with this role, we found that FOXO3a was dephosphorylated (Ser253) during unloading in both adult and old mouse soleus. Accordingly, the expression E3 ligases Atrogin-1 and Murf-1 were also elevated during atrophy. However, FOXO3a remained dephosphorylated and Atrogin-1 and Murf-1 tended to remain elevated in the old mice during recovery, compared to the adults (Figure 3B).

The ratio of LC3BI/II, an index of LC3B lipidation and autophagy activation, was elevated during unloading and reduced with recovery in adult and old mice (Figure 3C). Similar trends were seen for ATG7 and PINK which are proteins involved in mitophagy. PINK1, a protein thought to protect cells from stress-induced mitochondrial dysfunction, remained elevated during reloading in the old mice (Figure 3C). Taken together, these data show that the key regulators of protein synthesis and degradation respond as expected during unloading in both adult and old mice. However, following 3 days of reloading, there is persistent activation of protein degradation and inhibition of protein synthesis mediators in old mice.

**Figure 3.**
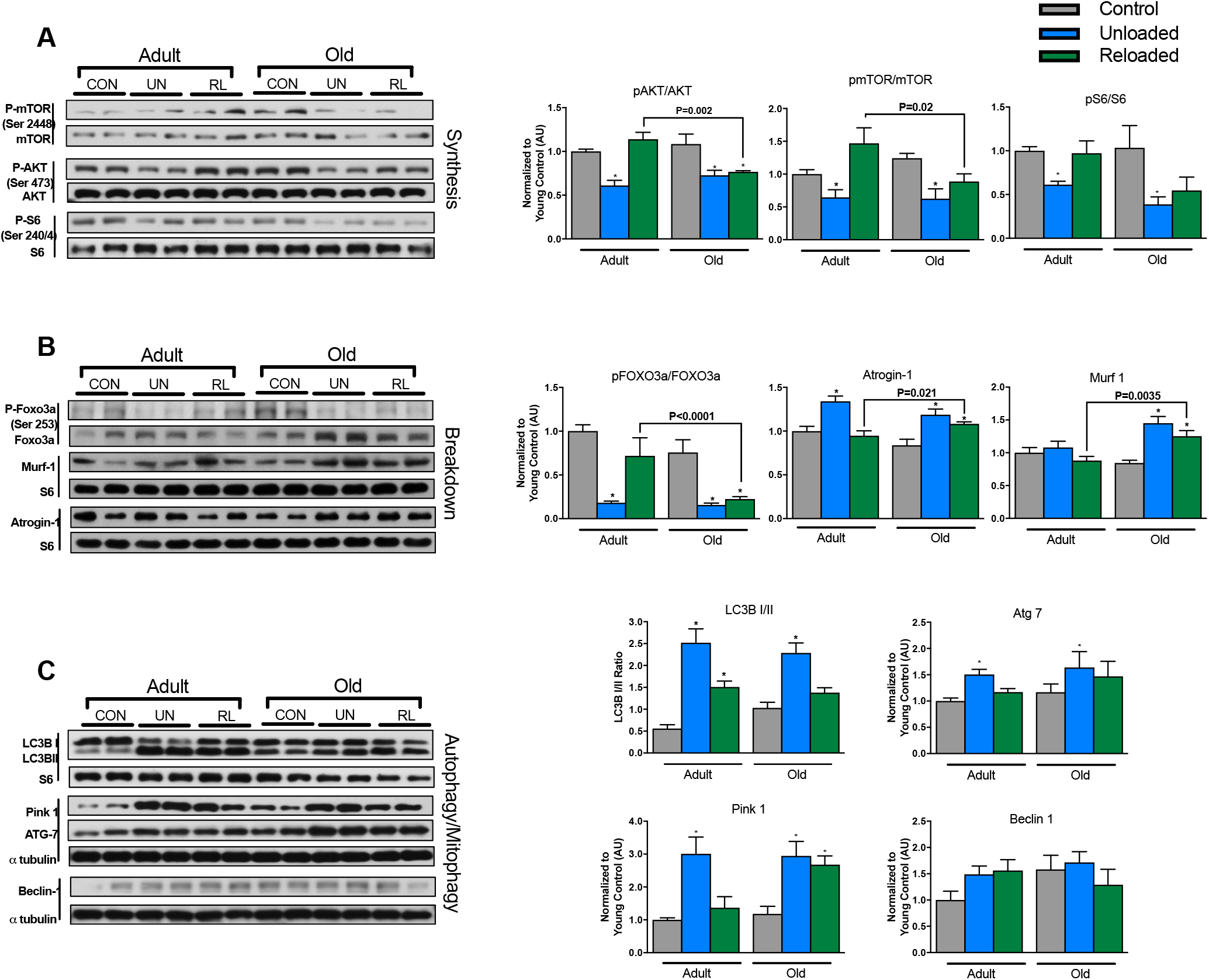
Mediators of protein synthesis are inhibited and breakdown are activated during unloading, a response that does not return to baseline following reloading in the old mice. <u>Panel A</u>, protein synthesis signaling: pAKT/AKT, pmTOR/mTOR, and pS6/S6 ratios. <u>Panel B</u>, protein breakdown signaling: pFOXO3a/FOXO3a ratio, Atrogin 1, and Murf 1 expression. <u>Panel C</u>, autophagy/mitophagy protiens: LC3B I/II ratio, ATG7, Pink 1, Beclin 1 expression. n = 5 per group. Data are presented as mean ± SEM. ** P <0.05 ANOVA/Bonferoni Correction. * P <0.05 v Control ANOVA/Bonferoni Correction.

### Mitochondrial function does not improve during recovery in old mice

Our central hypothesis is that alterations in mitochondrial energetics and fuel metabolism play a key role in the poor recovery of muscle mass in older animals. Functional assays of soleus mitochondria were conducted using saponin permeabilized myofiber bundles, an approach that preserves the native reticular structure of the mitochondria (Picard *et al*. 2011). When compared to adult, skeletal myofibers isolated from old control mice displayed lower respiration, OXPHOS content and calcium retention capacity (CRC), an index of apoptotic susceptibility, while H_2_O_2_ emission was elevated (Figure 4A-G). Following unloading, LEAK, complex I, I&II and FAO supported OXPHOS respiration and CRC in soleus were all reduced in both the adult and old mice. In addition, mitochondrial H_2_O_2_ emission was elevated in both young and aged animals (Figure 4A-F). Total cardiolipin and OXPHOS protein (measures of mitochondrial content) were both reduced with unloading in the adult mice, but remained unchanged in the older animals indicating that changes in function were independent of changes in mitochondrial content during unloading in the old animals (Figure 4G,H&J).

**Figure 4.**
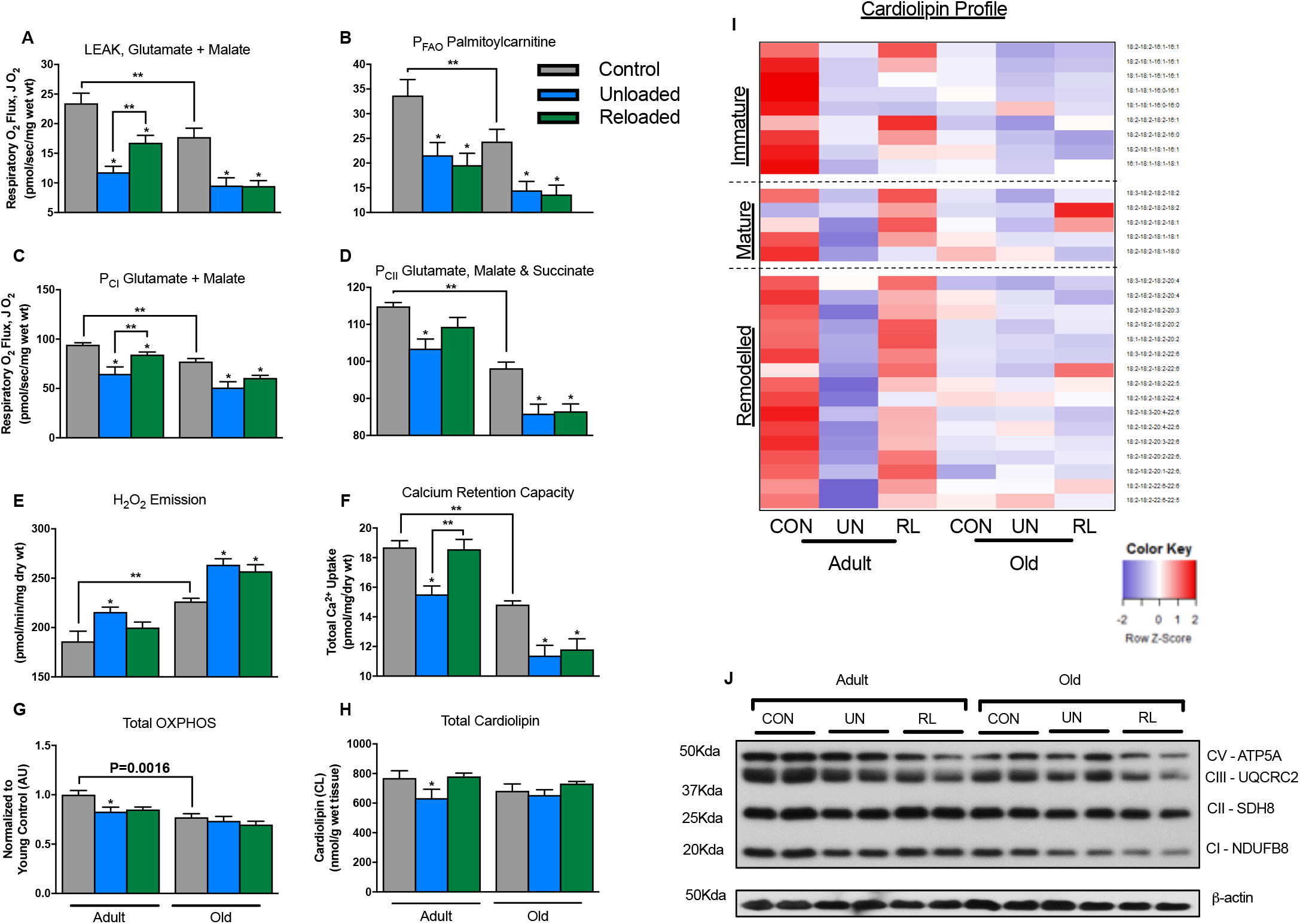
Mitochondrial function is impaired during unloading and improves during recovery only in adult mice. <u>Panel A</u>, Complex I supported LEAK (CI*L* or State 4) respiration. <u>Panel B</u>, Palmitoylcarnitine supported OXPHOS (State 3) respiration. <u>Panel C</u>, Complex I supported OXPHOS respiration. <u>Panel D</u>, Complex I&II supported OXPHOS respiration. n = 810 per group for respiration experiments. <u>Panel E</u>, Mitochondrial H_2_O_2_ emission. n = 8 – 10 per group. <u>Panel F</u>, Mitochondrial calcium retention capacity. n = 8 – 10 per group. <u>Panel G</u>, Total OXPHOS protein content by immunoblot. n = 5 per group. <u>Panel H</u>, Total cardiolipin content by tandem mass spectrometry. <u>Panel I</u>, heat map depicting change in individual immature, mature and remodeled cardiolipin species. n = 7 per group. <u>Panel J</u>, Representative immunoblot for OXPHOS proteins. Data are presented as mean ± SEM. ** P <0.05 ANOVA/Bonferoni Correction. * P <0.05 v Control ANOVA/Bonferoni Correction.

Following 3-days of recovery, complex I & II supported mitochondrial respiration, CRC, and H_2_O_2_ emission normalized in the young animals. Interestingly, FAO supported respiration did not recover in the young animals. Old mice did not recover any aspect of mitochondrial function as evidenced by sustained elevated H_2_O_2_ emission and suppressed CRC and respiration. Taken together, these data indicate that mitochondrial function does not improve during the early recovery phase in old mice.

We used shotgun lipidomics to quantify the cardiolipin pool (Han *et al*. 2006). Cardiolipin is a phospholipid specific to the inner mitochondrial membrane and critical for ETC integrity and mitochondrial function (Paradies *et al*. 2014). Cardiolipin species can be broadly categorized based on acyl chain length and degree of saturation, into: immature, mature and remodeled species. Immature cardiolipins are those that have heterogeneous acyl chain length and degree of saturation and typically contain one or more 16:1 acyl chains. Mature cardiolipins have more uniform acyl chain length and degree of saturation, for example, tetralinolyol cardiolipin (18:2-18:2-18:2-18:2). Remodeled cardiolipin species contain greater PUFA (typically 22:6) acyl chains and may have been remodeled due to oxidative insult (Chicco & Sparagna 2007). The soleus of older animals had slightly lower content of many cardiolipin species, again likely due to reduced mitochondrial content with aging (Figure 4I and Table 1S). Unloading elicited a striking reduction in many immature, mature and remodeled species of cardiolipin in adult mice. There was a partial recovery following reloading. The same dynamic response of the cardiolipin pool was not evident in the older mice.

### Divergent metabolomic response between adult and old mice during unloading and recovery

Targeted metabolomic profiling was conducted on soleus samples to quantify acylcarnitines, organic acids and amino acids in order to assess perturbations in mitochondrial fuel metabolism that may occur with unloading and reloading. We show for the first time a striking decrease in long-chain acylcarnitines during unloading in the adult animals (Figure 5A and Table 2S), in-line with the observations of others who have shown that contractile inactivity induces a switch to carbohydrate fuel source and reductions in fatty acid oxidation (Bergouignan *et al*. 2011). Interestingly, following reloading, acylcarnitines remained suppressed in the adult mice, consistent with lower FAO supported respiration. However, certain TCA cycle intermediates (fumarate, malate, and α-ketoglutarate) and amino acids (Gln, Glu, Ala, and Citrulline), increased during reloading in the adult soleus, consistent with increases in TCA/CI&II supported respiration (Figure 5B&C and Table 3&4S). These data suggest that FAO remains suppressed during the early reloading phase in adult muscle and that recovery of mitochondrial respiration is supported by glycolysis and TCA cycle anaplerosis via amino acids. The increase in C4, and C5 acylcarnitines during reloading are also consistent with increased amino acid catabolism and anaplerosis (Table 2S). In old mice, the metabolomic profile in response to unloading and reloading was distinctly different from that of the adult animals. The majority of long-chain acylcarnitines increased during unloading and reloading in the old mice (Figure 5A). This, in concert with reduced TCA cycle intermediates and reduced capacity for respiration, suggest that FAO outpaced TCA flux and resulted in accumulation of acylcarnitine species. Interestingly, many of the amino acids that were increase upon reloading in the adult animals did not rise in the older animals (Figure 5C).

**Figure 5.**
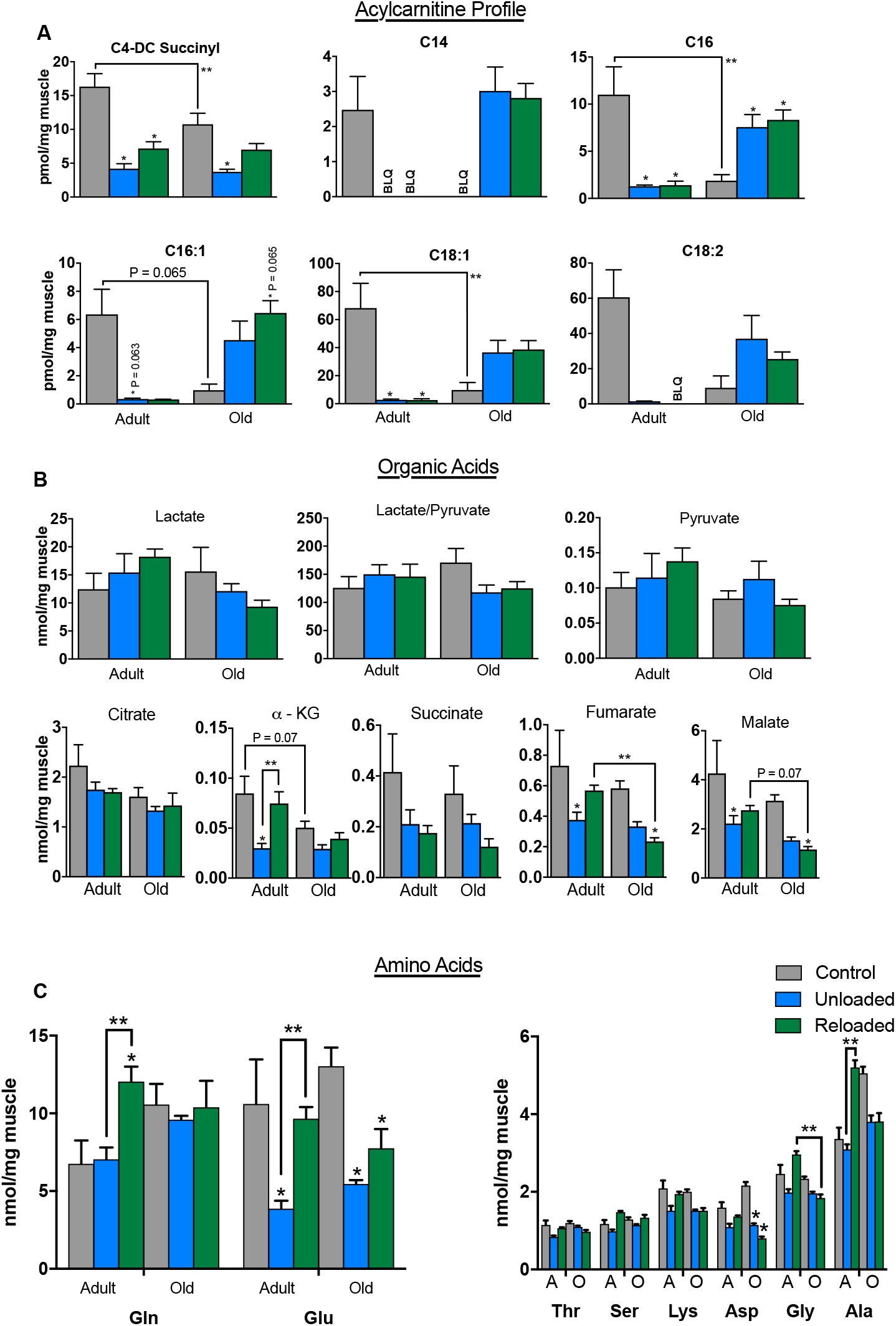
Acylcarnitine, organic acid and amino acid profiles during unloading and reloading. <u>Panel A</u>, Acylcarnitine profile in soleus of adult and old mice during control, unloading and reloading conditions. Acyl chain length and degree of saturation are denoted by the numbers. <u>Panel B</u>, metabolite profiles showing lactate and pyruvate and organic acid intermediates of the tricarboxylic acid (TCA) cycle. <u>Panel C</u>, amino acids in soleus of adult and old mice during control, unloading and reloading conditions. BLQ = below limit of quantification. n = 6 per group. Data are presented as mean ± SEM. ** P <0.05 ANOVA/Bonferoni Correction. * P <0.05 v Control ANOVA/Bonferoni Correction.

### Transcriptomic response to unloading and reloading is similar in adult and old mice

Gene expression profiling was conducted on soleus samples from the 6 groups using RNA sequencing. Our goal was to determine whether the divergent mitochondrial energetic and fuel metabolic responses to unloading and particularly reloading in old mice were underpinned by a distinct transcriptomic response. A summary of the number of differentially regulated genes is presented in Table 7S. The database for annotation, visualization and integrated discovery (DAVID) v6.8 was used to identify enriched biological themes via functional annotation (Dennis *et al*. 2003). The top significantly up- and down-regulated functional category terms for each group comparison are presented in Supplemental Tables 8-11S. Terms related to mitochondrial function and fuel metabolism were downregulated in during unloading for both adult and old mice (Table 8&9S). Surprisingly, mitochondria related terms were also significantly downregulated during reloading for adult and old mice (Table 10&11S). These findings were confirmed by qRT-PCR assay for expression of key nuclear transcription factors that drive mitochondrial energy metabolism, including: *Ppargc1a* (PGC-1α), *Esrrg* (ERRγ), and *Esrra* (ERRα) (Figure 6A).

**Figure 6.**
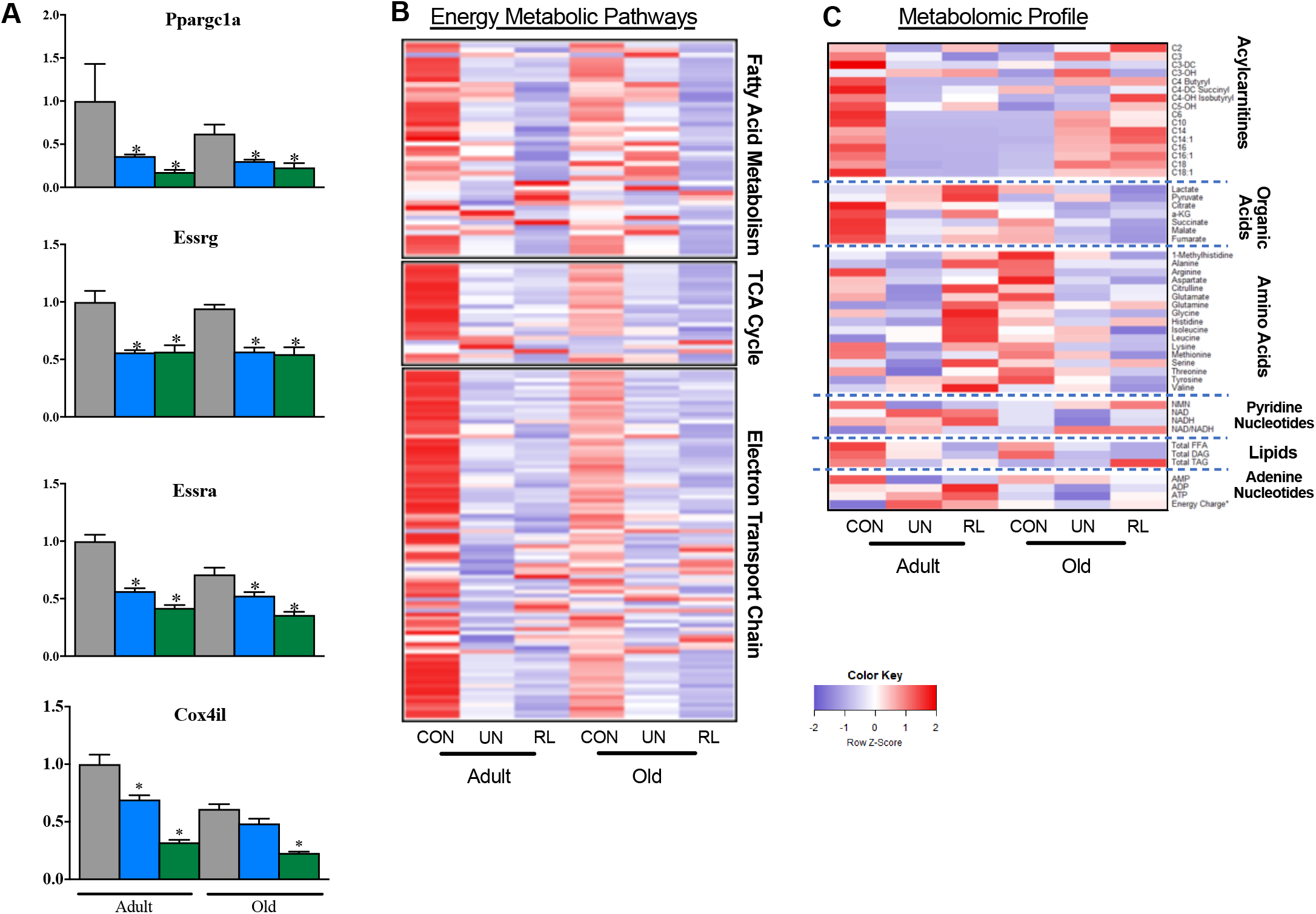
Comparative analysis of transcriptomic and metabolomics profiles during unloading and reloading. <u>Panel A</u>, qRT-PCR assay for expression of key nuclear transcription factors that drive mitochondrial energy metabolism. <u>Panel B</u>, heat map containing RNA sequencing data set representing the level of expression of genes with reads per kilobase per million mapped reads (RPKM) >1, plotted as a centralized Z-score for each gene in fatty acid metabolism, tricarboxylic acid (TCA) cycle, and electron transport chain/oxidative phosphorylation pathways defined by Ingenuity Pathway analysis. <u>Panel C</u>, heat map containing the levels of acylcarnitines, organic acids, amino acids, pyridine and adenine nucleotides and total level of free fatty acids (FFA), diacylglycerol (DAG) and triacylglycerol (TAG). For all heat maps, relative downregulation is in blue and upregulation in red. The intensity of color indicates the magnitude of the change.

We next compared the collective results of the transcriptomic and metabolomic profiling. This comparative analysis is depicted by heat maps (Figure 6B&C) which display the direction of differential expression. The analysis was restricted to a subset of genes involved in mitochondrial fuel and energy metabolism relevant to the metabolites measured in the targeted metabolomics profile. This comparison highlights striking differences in the metabolomic and transcriptomic responses during the reloading condition for adult and old mice. In adult mice, the recovery of mitochondrial respiratory capacity occurs along with partial normalization of metabolites, particularly amino acids and organic acids, but not acylcarnitines. In contrast, FAO, TCA and ETC genes remained repressed. In old mice, the metabolite response during recovery is distinctly different. Namely, acylcarnitines were elevated while organic acids and amino acids remained low. Moreover, FAO, TCA and ETC genes remained repressed. Taken together, these results indicate that mitochondrial energetics gene and metabolite profiles are repressed in concert with unloading and muscle atrophy. However, metabolite levels clearly distinguish the early muscle hypertrophy response in adult animals versus the lack of recovery in old animals upon reloading and to a far greater degree than mitochondrial energetics gene profiles.

## DISCUSSION

During aging, skeletal muscle exhibits a progressive loss of mass and strength known as sarcopenia (Rosenberg 1989), and an attenuated ability to recover muscle mass following acute periods of disuse or unloading. Poor recovery of muscle mass following periods of disuse is a significant health concern for older adults that leads to lower activities of daily living and increased risk of frailty and mobility disability. The mechanisms responsible for the lack of muscle mass recovery in old mice following disuse have not been fully elucidated but are likely complex and multifactorial. Investigations to date have focused on attenuated protein synthesis possibly due to ER stress and impaired neuromuscular transmission (Baehr *et al*. 2016). In separate lines of investigation, aging has also been linked with insulin resistance and mitochondrial dysfunction in skeletal muscle (Konopka & Sreekumaran Nair 2013). Here, we report that 22-24 month old mice with low muscle mass and insulin resistance, display poor early recovery of muscle mass following 10 days of hind limb unloading. We took an unbiased approach to identify changes in energy metabolic gene expression and metabolite pools and show for the first time that persistent mitochondrial dysfunction and a dysregulated metabolic response underlie poor early recovery of muscle mass in old mice. Importantly, this is linked to worsened whole-body insulin resistance. Our results demonstrate the power of using unbiased molecular profiling to define signatures of complex physiological states combined with deep phenotyping of mitochondrial energetics.

Physical inactivity results in attenuated mitochondria capacity along with a shift in fuel selection to carbohydrate and suppressed FAO (Bergouignan *et al*. 2011). Here, in adult mice, we show a distinct remodeling of soleus mitochondrial energetics and fuel metabolism during unloading as evidenced by reduced respiration, elevated H_2_O_2_ emission and reduced long-chain acylcarnitines and organic acids. This change in mitochondrial phenotype occurred along with striking reductions in many individual molecular species of cardiolipin. Indeed, synthesis and remodeling of cardiolipin is crucial for the structural and functional integrity of the mitochondria. Our findings expand upon the work of Ostojic et al., others who reported decreased cardiolipin during muscle atrophy (Ostojic *et al*. 2013).

Insulin signaling intersects with protein synthesis and degradation pathways via AKT/FOXO/mTOR. Hence, a logical supposition is that baseline insulin resistance worsens muscle atrophy response during disuse. Similarly, mitochondrial dysfunction and ROS emission are linked to activation of protein degradation pathways and may worsen muscle atrophy due to disuse (Romanello & Sandri 2010). Indeed, older adults with type 2 diabetes, a condition defined by muscle insulin resistance and mitochondrial dysfunction, have an accelerated loss of muscle mass over time (Park *et al*. 2009), indicating a propensity for catabolic metabolism. However, we found that insulin resistant old mice with mitochondrial dysfunction did not have an exacerbated muscle atrophy during disuse. The relative magnitude of muscle loss and alterations in the activity (phosphorylation state) of AKT/mTOR/FOXO3a, and the majority of endpoint measurements were similar in adult and old animals in response to unloading. These findings suggest that the presence of insulin resistance or mitochondrial dysfunction in aging does not influence the muscle atrophy response due to disuse or unloading.

Although impaired recovery by aged muscle is reasonably well documented, the mechanisms responsible for the lack of growth are not clearly defined. We show for the first time that impaired early recovery of muscle mass may be exacerbated by insulin resistance and persistent mitochondrial dysfunction in old animals. In stark contrast to adult mice, there was no change in the cardiolipin species of old mice that likely contributed to persistent mitochondrial dysfunction. Cardiolipin synthesis and remodeling is critical for correct cristae folding and functional integrity of the ETC complexes. The increase in acylcarnitines (long chain in particular) in concert with reductions in particular TCA cycle organic acids (fumarate and malate), and no change in pyruvate or lactate, suggest a bottleneck in carbon flux from ß-oxidation into the TCA cycle which could reduce the capacity to generate the reducing equivalents needed to produce ATP contributing to the anabolic response. The change in muscle metabolite pools during reloading in old mice is similar to that observed with high fat feeding and obesity (Koves *et al*. 2008). In addition, the metabolite profile served to robustly distinguish the phenotype of poor muscle recovery in old mice. Our findings are also in line with observations that insulin resistance, obesity, and elevated plasma free fatty acids are linked with blunted amino acid stimulated muscle protein synthesis (Murton *et al*. 2015; Stephens *et al*. 2015). Further studies are needed to examine whether alterations in mitochondrial fatty acid import or FAO may impact early recovery of muscle in old mice, potentially via improved insulin sensitivity.

Interestingly, while metabolite profile distinguished the phenotype of poor muscle recovery in old mice, transcriptomic profiles for energy metabolic pathways were similar between old and adult mice during unloading and reloading. In addition, the recovery of mitochondrial respiratory capacity in adult mice occurred along with partial recovery of metabolites, while FAO, TCA and ETC genes remain repressed. Indeed, FAO genes appeared to be further repressed during unloading in adult mice. These data suggest that the early recovery of mitochondrial function following reloading is, seems to be independent of FAO and mediated by cardiolipin remodeling and alterations in metabolite profile. This, in turn, could affect post translational modification and activity of ETC, although this remains to be tested. Taken together, these results indicate that mitochondrial energetics, gene and metabolite profiles are repressed in concert with unloading and muscle atrophy.

In summary, our data suggest that dysregulated FAO and elevated H_2_O_2_ emission from mitochondria likely contribute to the lack of recovery of soleus muscle mass following unloading in sarcopenic mice. These findings suggest that pathways governing muscle fuel metabolism and mitochondrial energetics might harbor promising therapeutic targets for improving the recovery of muscle mass following periods of disuse in older patients.

## EXPERIMENTAL PROCEDURES

### Animals

All animal experiments and euthanasia protocols were conducted in strict accordance with the National Institute of Health guidelines for humane treatment of animals and approved by the Institutional Animal Care and Use Committee at the Sanford-Burnham Medical Research Institute at Lake Nona. 6-month and 22-24 month old male C57BL/6J mice were obtained from the National Institute on Aging rodent colony (Charles River Laboratories, Madison, WI). Young and aged animals were divided into the following experimental groups: 10-day of hind limb unloading (UN), 10-day of hind limb unloading followed by 3 days of reloading recovery (RL), and control (CON). Mice were fed ad libitum with a standard chow diet (#2916, Harlan-Teklad, Houston, Tx) and water/hydrogel (ClearH2O, Westbrook, ME, US) and housed at 22C with a 12h light/dark cycle.

### Tail suspension hind limb unloading

Tail suspension is the one of the most commonly used animal models of musculoskeletal non-weight bearing. Prior to tail suspension experiments the individual mice acclimated in single cages and routine handling for 3 days. Tail suspension was performed using a modified version of the Morey-Holton and Globus protocol (Morey-Holton & Globus 2002). Briefly, a small metallic hook is taped to the base of the tail using non-abrasive adhesive tape wrapped in a helical pattern. The hook is then attached to a small swivel key chain that is attached to a metal rod that runs the length of the microisolator cage. The mice could move on a y-axis and rotate 360 degrees and so had access to all areas of the cage. The hind limbs are maintained just off the cage floor with the body of the mouse at ~30^o^ angle from the cage floor. The mice can move freely and the angle and height of the mice are checked daily. The control mice were separated into the individual cages with the exactly the same conditions as unloading groups, but without tail suspension.

After 10 days of UN, RL, or CON, the mice were fasted for 4 hours and sacrificed by CO2 asphyxiation and cervical dislocation, or anesthetized with sodium pentobarbital (IP administration, 50mg/kg) for muscle harvested for metabolomics analysis. The soleus, gastrocnemius, and quadriceps femoris muscle groups were immediately harvested and weighed. The soleus muscle from right hind limb was used for fresh tissue assay immediately and other muscles were snap-frozen in liquid nitrogen for other analysis.

### Cardiometabolic Phenotyping

<u>*Body Composition*</u>. Measurement of fat and lean mass (g) is accomplished using a LF90II TD-NMR (Bruker, Madison, WI). Conscious mice are placed in a plastic restraint tube that is inserted into the instrument. Measurements are obtained in less than 1 minute. <u>*Ambulatory cage activity*</u>. Locomotor activity measurements are obtained using a Promethion Mouse Multiplexed Metabolic System (Sable Systems International, Las Vegas, NV). Body mass is collected using a housing cubby connected to a precision scale. Locomotor activity along the X, Y and Z planes is measured using infrared beams that span the home cage. Data are collected by a host computer. <u>*Insulin-stimulated Tissue Glucose Uptake*</u>, was assessed as described in the Supplemental Information section. Processing of tissues for glucose uptake was determined as previously described (Ayala *et al*. 2007), and as detailed in the Supplemental Information section.

### Myofiber Bundle Preparation

Permeabilized fiber bundles (1-3mg) were prepared immediately following tissue harvest, as previously described (Coen *et al*. 2015) and as detailed in the Supplemental Information section.

### Mitochondrial respiration

Respirometry assays were conducted using an Oxygraph-2k (Oroboros Instruments, Innsbruck, Austria). The myofiber bundles were gently placed into the respirometer chambers and after a stable baseline was reached, the assay protocol was run in duplicate at 37^o^C and between 350-200 nmol of O_2_ in Buffer Z with blebbistatin (25μM). The assay protocols are described in the Supplemental Information section.

### Mitochondrial H_2_O_2_ Emission

H_2_O_2_ emission was measured with Amplex Red reagent which reacts with H_2_O_2_ to produce the stable fluorescent compound resorufin. Resorufin fluorescence was monitored using a Fluorescence Spectrometer (Perkin Elmer LS50B, Waltham, Massachusetts, USA) with temperature control (37^o^C) and magnetic stirring at excitation/emission 563/587nm. The assay was run with buffer Z containing 5000U/ml CuZn-SOD, 25μM blebbistatin, 50μM Amplex Ultra-Red, and 6U/ml horseradish peroxidase. Briefly, the muscle fiber bundles were added to the reaction buffer with 10ug/ml oligmycin, 10mM Glutmate, 2mM Malate and 10mM Succinate. The rate of emission of H_2_O_2_ (pmol) was calculated from previously established fluorescence intensity standard curves with known concentrations of H_2_O_2_, after correcting for the rate of change in background fluorescence. Following each experiment, myofiber bundles were dried and weighed. Rate of H_2_O_2_ emission was expressed as pmol/min/mg dry weight.

### Mitochondrial calcium retention capacity

A continuous, spectrophotometric assay was utilized to measure mitochondrial calcium retention capacity within soleus fiber bundles, as detailed in the Supplemental Information section.

### RNA Sequencing and Informatics

Details of sample preparation, library preparation sequencing, mapping and differential gene expression analysis are described in the Supplemental Information section. <u>*Heat maps*</u>. The normalized (z-score) mean of each cohort was visualized in the heatmap, created with the “heatmap.2” function in the “gplots” package within the statistical program R (<u>http://www.r-project.org/</u>). <u>*Gene Ontology (GO) annotation*</u>. The database for annotation, visualization and integrated discovery (DAVID) v6.8 was used to identify enriched biological themes via gene ontology (GO) annotation. The gene expression data discussed in this publication have been deposited in NCBI’s Gene Expression Omnibus and are accessible through GEO Series accession number GSE102284.

### Real-time quantitative-PCR

Details of sample preparation and gene expression analysis by PCR are described in the Supplemental Information section.

### Western Blot

Frozen soleus was prepared for immunoblot as described previous (Lee *et al*. 2017) and as detailed in the Supplemental Information section.

### Lipidomics Analysis

Multi-dimensional mass spectrometry-based shotgun lipidomics (MDMS-SL) was employed to measure and characterize the lipid patterns in mouse soleus muscle. The analytical procedures for the lipidomics analysis are detailed in the Supplemental Information section.

### Metabolomics Analysis

Acylcarnitines, organic acids and amino acids were quantified using internal calibration standards and mass spectrometry as described in detail in the Supplemental Information section.

### Statistical Analysis

All data are represented as the mean ± SEM. All the statistical analyses were performed by GraphPad Prism. The differences between groups were conducted using ANOVA or *t* -test (paired and unpaired) approaches with Tukey’s/Bonferroni Multiple Comparison Test. when appropriate p <0.05, it was considered significant.

## ACKNOWLEDGEMENTS/FUNDING

This study was supported by funding from the National Institutes of Health | National Institute on Aging (K01 AG044437) awarded to PMC. We are grateful for the excellent technical assistance of the staff at the vivarium, the Cardiometabolic Phenotyping core and the Metabolomics core of the Sanford Burnham Prebys Medical Discovery Institute at Lake Nona. We are also grateful to Feng Qi for bioinformatics support and Fanchao Yi for statistics support. The authors have no conflicts of interest to declare.

## AUTHORS CONTRIBUTIONS

XZ, MBT, TCL, DPK, XH, SJG, JA, RBV, PMC performed experiments, and analyzed and interpreted the data. XZ and RBV contributed to data interpretation and writing the manuscript. All co-authors reviewed and approved the manuscript. PMC contributed to the study concept and design, statistical analysis, interpretation of the data and wrote the manuscript. PMC is the guarantor of the data.

